# Representation of navigational affordances and ego-motion in the occipital place area

**DOI:** 10.1101/2024.04.30.591964

**Authors:** Frederik S. Kamps, Emily M. Chen, Nancy Kanwisher, Rebecca Saxe

## Abstract

Humans effortlessly use vision to plan and guide navigation through the local environment, or “scene”. A network of three cortical regions responds selectively to visual scene information, including the occipital (OPA), parahippocampal (PPA), and medial place areas (MPA) – but how this network supports visually-guided navigation is unclear. Recent evidence suggests that one region in particular, the OPA, supports visual representations for navigation, while PPA and MPA support other aspects of scene processing. However, most previous studies tested only static scene images, which lack the dynamic experience of navigating through scenes. We used dynamic movie stimuli to test whether OPA, PPA, and MPA represent two critical kinds of navigationally-relevant information: navigational affordances (e.g., can I walk to the left, right, or both?) and ego-motion (e.g., am I walking forward or backward? turning left or right?). We found that OPA is sensitive to both affordances and ego-motion, as well as the conflict between these cues – e.g., turning toward versus away from an open doorway. These effects were significantly weaker or absent in PPA and MPA. Responses in OPA were also dissociable from those in early visual cortex, consistent with the idea that OPA responses are not merely explained by lower-level visual features. OPA responses to affordances and ego-motion were stronger in the contralateral than ipsilateral visual field, suggesting that OPA encodes navigationally relevant information within an egocentric reference frame. Taken together, these results support the hypothesis that OPA contains visual representations that are useful for planning and guiding navigation through scenes.

## Introduction

Humans expertly use vision to plan and guide navigation through scenes. The past several decades have revealed a network of at least three cortical regions that respond selectively to visual scene information, including the occipital place area (OPA) (Dilks et al., 2013), parahippocampal place area (PPA) (Epstein and Kanwisher, 1998), and medial place area (MPA; also known as retrosplenial complex) (Maguire, 2001; Silson et al., 2016). It has recently been hypothesized that these regions are functionally dissociated, with PPA supporting scene categorization (e.g., am I in a kitchen or a forest?), MPA supporting memory-guided navigation (e.g., which way should I head to find my hotel, six blocks away?), and OPA supporting visually-guided navigation (e.g., can I walk to the left or the right?) (Dilks, Kamps, and Persichetti, 2022).

Of the three scene-selective regions, the least is known about OPA, which is small and variably located across adults (Julian et al., 2012). Nevertheless, growing evidence supports the hypothesis that OPA is specialized for visually-guided navigation (Dilks et al, 2011; Persichetti and Dilks, 2016; Kamps et al., 2016a). For example, OPA responds significantly more when participants are asked to indicate how they would navigate through a scene than when asked to indicate the category of the same scene (PPA showed the opposite pattern, responding more to the categorization task than the navigation task; MPA showed no preference) (Persichetti and Dilks, 2017). Two further studies found that OPA (but not PPA or MPA) is sensitive to navigational affordances: for example, patterns of activation in OPA, in response to static snapshots of novel scenes, contain reliable information about the location of open doorways (Bonner and Epstein, 2017), and the presence of spatial boundaries that limit locomotion (Park and Park, 2020). OPA is also sensitive to the perceived distance (Persichetti and Dilks, 2016) and direction (Dilks et al., 2011) of scene information, suggesting an ego-centric frame of reference, which is critical for planning and guiding one’s own movements through space.

Most studies to date (including all studies cited above) have relied on static images to study OPA responses. Yet in real life, navigation is inherently dynamic: visual scenes are seen in motion as the observer moves through them (i.e. optic flow). Accordingly, if OPA is specialized for visually-guided navigation, then the full functional profile of OPA may only be revealed during dynamic, first-person motion through scenes. Consistent with this possibility, five studies suggest that OPA is sensitive to dynamic scene information. Kamps et al. (2016b; 2020) and Suzuki, Kamps et al. (2021) showed that OPA responds significantly more to dynamic scenes depicting first-person perspective motion through scenes than to static images taken from these same scene videos (with no such motion enhancement found for faces or objects). Similarly, Hacialihafiz et al. (2016) showed that OPA responds significantly more to dynamic scene displays depicting linear horizontal motion (e.g., panning across a scene image from left to right) than static versions of those same displays. Finally, Sulpizio et al. (2020) found that OPA responds more to point-light displays depicting ego-motion-compatible optic flow than random motion. For all five studies, responses to motion information in OPA were greater than those in PPA or MPA, providing initial evidence for a functional dissociation in the scene network based on motion sensitivity. However, none of these studies has explored the finer grained information that OPA extracts from dynamic scenes. It is therefore unclear to what extent OPA responses to dynamic scenes reflect information processing relevant to planning and guiding navigation.

To better understand the nature of dynamic scene representations in OPA, we tested whether OPA extracts two kinds of navigationally-relevant information in dynamic scenes: 1) *navigational affordances* (is there an open doorway to the left, right, or both?) and 2) *ego-motion directions* (moving forward or backward, turning left or right), as well as the conflict of these two kinds of information (e.g., turning toward versus away from an open doorway) (Figure 1). We predicted that OPA represents both navigational affordances and ego-motion directions, as well as their conflict, and that responses to such navigationally relevant information would be stronger in OPA than both PPA and MPA, consistent with the hypothesized functional dissociations between these regions. An important further question concerns the extent to which responses to navigationally-relevant information in OPA could reflect low-level, retinotopic properties of the stimuli that are confounded with navigationally relevant information (e.g., left turn stimuli will generate greater motion energy in the right visual field, compared with the left visual field), rather than more abstract, higher-level navigationally-relevant invariances (Groen et al., 2017). To address this question, we also compared responses in OPA with those in early visual cortex (EVC), which has robust retinotopic spatial structure and is highly sensitive to motion energy, providing a proxy for lower-level, retinotopic processing. If OPA represents higher-level visual information relevant for navigation, then responses to navigationally relevant information in OPA will differ from those in EVC. The results confirmed both sets of predictions. Note though that the analyses reported here include minor deviations from preregistered analysis plan. Results from the originally planned analyses are described in the supplemental materials.

**Figure 1.**
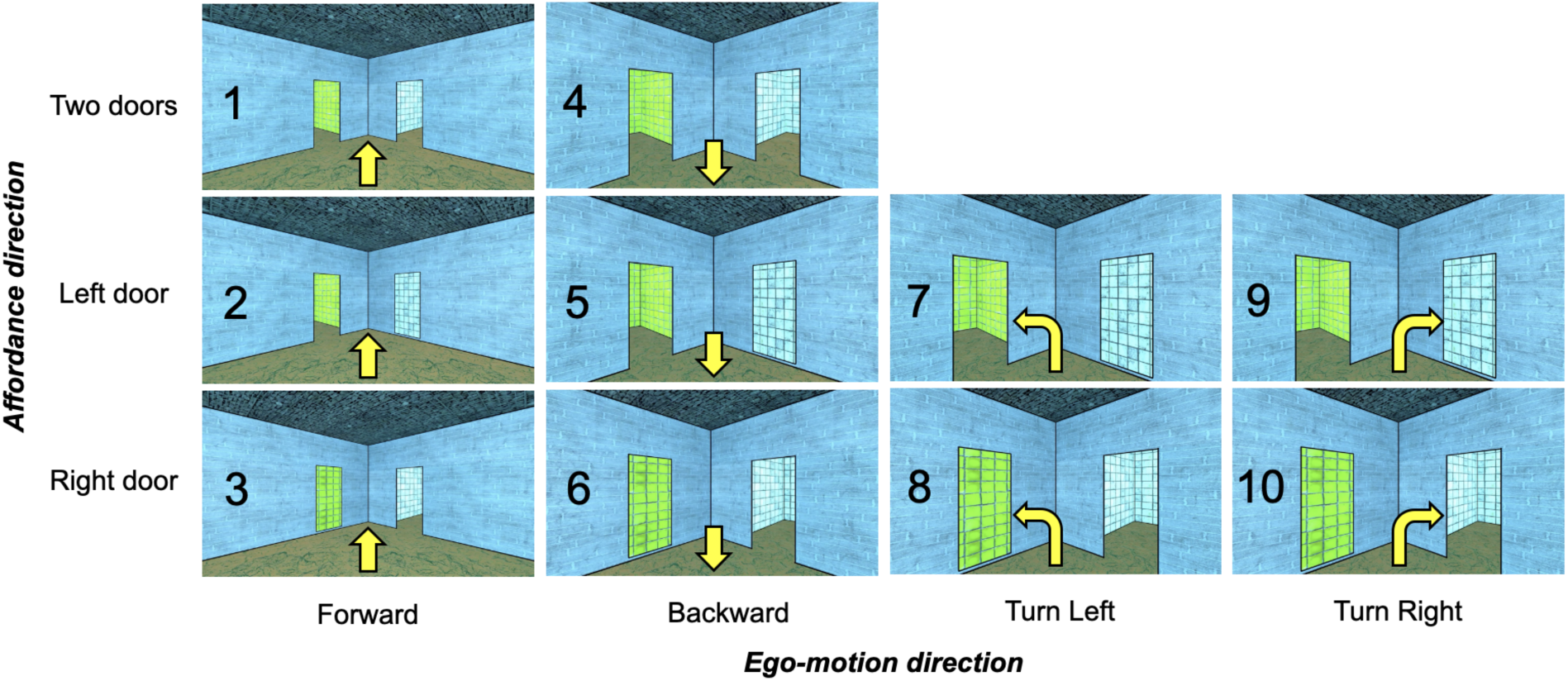
Dynamic movie stimuli. Adult participants viewed 3s video clips in an event-related design. The still images shown here depict the first frame of each movie stimulus. Condition numbers are listed over each stimulus. The direction of navigational affordance information (i.e., open doorways) in each scene is organized by row, depicting either two affordances (top row), an affordance to the left, but not right (middle row), or an affordance to the right, but not the left (bottom row). The direction of ego-motion is organized by column, and further indicated by the yellow arrow over each image, depicting either forward (first column), backward (second column), left turn (third column), or right turn (fourth column) ego-motion.

## Results

### OPA represents navigational affordances in dynamic scenes

If OPA represents navigational affordances in dynamic scenes, then OPA responses should differ depending on the location of navigational affordances present in the scene. To test this prediction, we analyzed conditions depicting ego-motion through scenes with either i) an open doorway to the left and a painting on the right (conditions 2 and 5), ii) an open doorway to the right and a painting on the left (conditions 3 and 6), or iii) two open doorways (conditions 1 and 4). We explored responses to these conditions in a series of multivariate and univariate analyses. Conditions with turns (i.e., conditions 7-10) were not included in these analyses, since turning changes the direction of navigational affordance information during the 2.5-second trial.

For multivariate analyses, we predicted more similar responses (measured using Euclidean distance) between each of the three affordance conditions (door to the left, right, or both) and itself (across split halves of the data; “within” conditions) than between each affordance condition and the others (also calculated across split halves of the data; “between” conditions). Full hypothesis matrices are shown in Figure 2a. Further, we predicted stronger decoding of navigational affordance information in OPA than PPA and MPA. We first tested this prediction using all possible pairs of conditions (i.e., the “overall” matrix in Figure 2a). Indeed, paired samples t-tests (one tailed) revealed significant navigational affordance decoding in OPA (t_(14)_ = 3.34, p = 0.002), a smaller but significant effect in PPA (t_(14)_ = 2.15, p = 0.03), and no significant effect in MPA (t_(14)_ = 1.28, p = 0.11) (Figure 2b). Directly comparing between regions, a 3 (region: OPA, PPA, MPA) x 2 (affordance: within, between) repeated measures ANOVA revealed a significant region x affordance interaction (F_(1.21, 15.74)_ = 5.00, p = 0.04), driven by stronger responses to affordance information in OPA than both PPA (post-hoc interaction contrast; F_(1,13)_ = 5.01, p = 0.04) and MPA (post-hoc interaction contrast; F_(1,13)_ = 5.51, p = 0.04). Next, to provide a stronger test of navigational affordance representation, we tested for representation of navigational affordances generalizing over ego-motion information (moving forward versus moving backward) by limiting comparisons to pairs of conditions that differed in ego-motion information, thereby minimizing the lower-level visual similarity of the “within” condition comparisons. Paired samples t-tests (one tailed) again revealed significant navigational affordance decoding in OPA (t_(14)_ = 3.43, p = 0.002), a smaller but significant effect in PPA (t_(14)_ = 2.80, p = 0.007), and no significant effect in MPA (t_(14)_ = 0.22, p = 0.42) (Figure 2b). Directly comparing between regions, a 3 (region: OPA, PPA, MPA) x 2 (affordance: within, between) repeated measures ANOVA revealed a significant region x affordance interaction (F_(2,26)_ = 6.22, p < 0.001), driven by stronger responses to affordance information in OPA than both PPA (post-hoc interaction contrast; F_(1,13)_ = 6.21, p = 0.03) and MPA (post-hoc interaction contrast; F_(1,13)_ = 7.54, p = 0.02).

**Figure 2.**
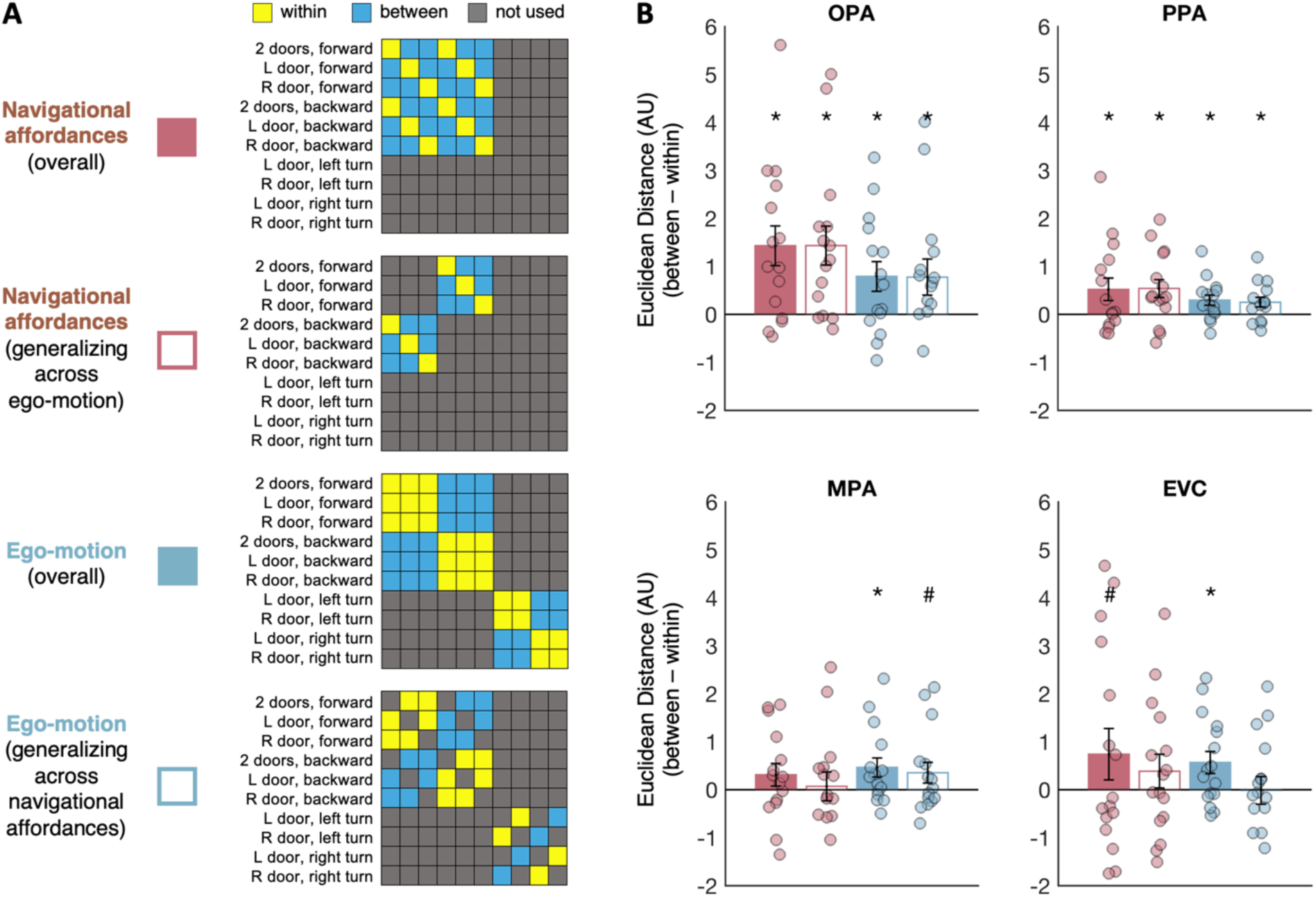
Multivariate analyses of navigational affordance and ego-motion representation. **A)** Hypothesis matrices used to test navigational affordance and ego-motion representation. If OPA represents navigational affordances (top matrix) and/or ego-motion (second matrix from bottom), then voxelwise patterns of activity will be more similar (i.e., smaller Euclidean distance) between conditions with the same navigational affordance or ego-motion information (yellow cells) than conditions with different information (blue cells). Navigational affordance and ego-motion information were tested “overall” (solid bars), as well as generalizing across the other kind of information (empty bars; e.g., testing navigational affordance only across conditions that differ in ego-motion, and vice versa for ego-motion information, generalizing across navigational affordances). **B)** Results of the four multivariate decoding analyses in the occipital place area (OPA), parahippocampal place area (PPA), medial place area (MPA), and an early visual cortex (EVC) control region. Bar plots indicate the difference score for the “between” conditions minus the “within” conditions; responses greater than zero indicate more similar responses (i.e., smaller Euclidean distance) for the “within” than “between” condition pairs, and thus significant decoding. AU indicates arbitrary units of Euclidean distance in fMRI response space. Error bars represent the standard error of the mean. Markers depict data from individual participants.

In univariate analyses, we tested the prediction that OPA will respond more to scene features that afford navigation (i.e., open doorways) versus those that do not (i.e., paintings). To test this prediction, we focused on the single-door scene conditions with either i) an open doorway to the left and a painting on the right (conditions 2 and 5), or ii) an open doorway to the right and a painting to the left (conditions 3 and 6). Given previous evidence that OPA shows a strong contralateral visual field bias (MacEvoy and Epstein, 2007; Silson et al., 2015; Silson et al., 2022), we predicted that the OPA in each hemisphere would respond more to doors than paintings in the contralateral visual field (i.e., right OPA responding more to doors in the left visual field than the right visual field, and vice versa for left OPA). To maximize statistical power, responses to contralateral doors vs. paintings were averaged across hemispheres, and then compared using a paired samples *t*-test (one-tailed). We found significantly stronger responses to contralateral doors than paintings in OPA (t_(14)_ = 3.95, p < 0.001), as well as marginal effects in PPA (t_(14)_ = 1.73, p = 0.05) and MPA (t_(14)_ = 1.57, p = 0.07) (Figure 3a). Directly comparing between regions, a 3 (region: OPA, PPA, MPA) x 2 (affordance: contralateral, ipsilateral) repeated measures ANOVA revealed a significant region x affordance interaction (F_(1.14, 15.93)_ = 9.28, p = 0.006), with a stronger contralateral affordance preference in OPA than both PPA (post hoc interaction contrast; F_(1,14)_ = 10.79, p = 0.005) and MPA (post hoc interaction contrast; F_(1,14)_ = 8.92, p = 0.01).

**Figure 3.**
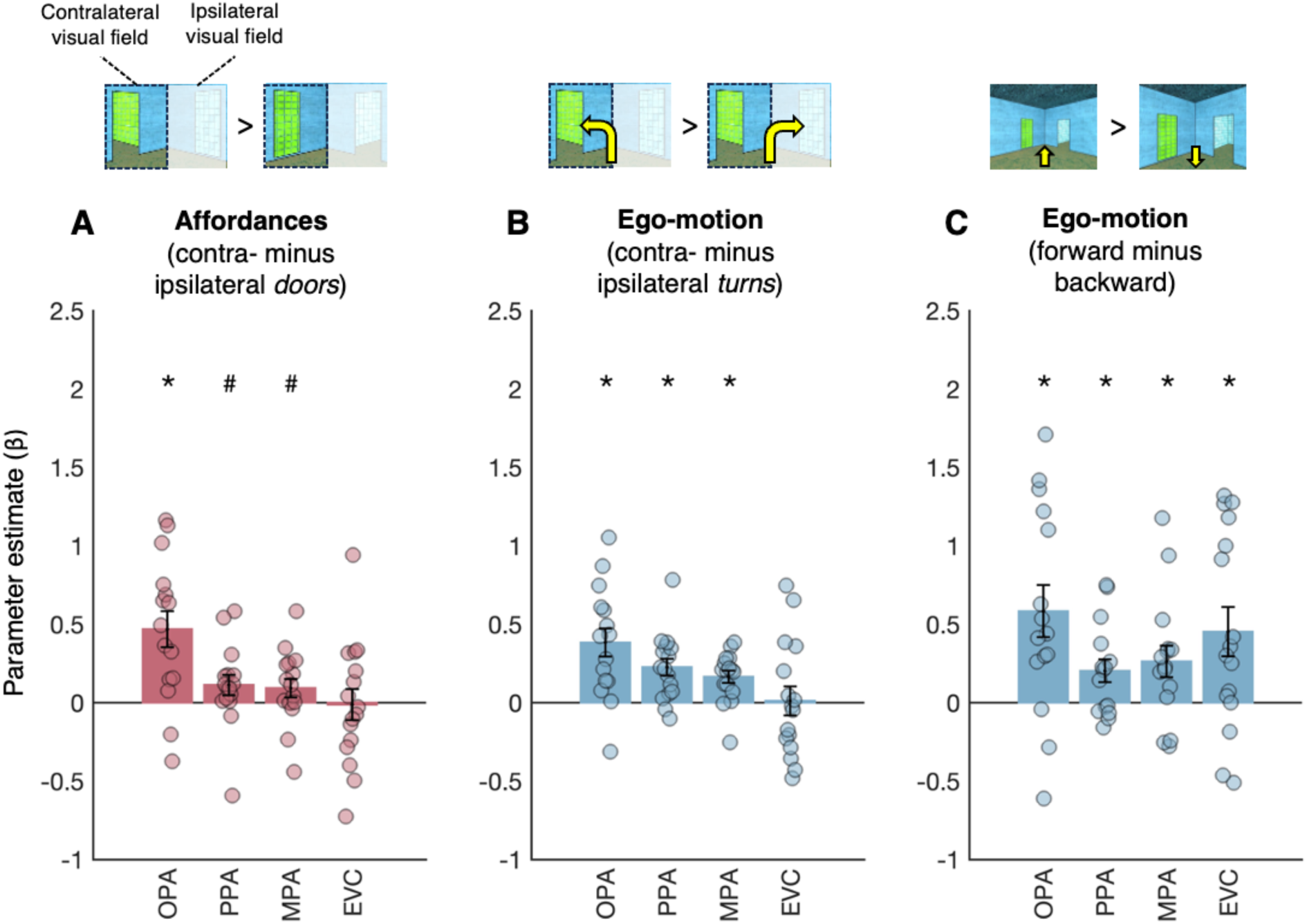
Univariate analyses of navigational affordance and ego-motion representation. A) Navigational affordance representation was tested based on the visual field (contralateral vs. ipsilateral) in which the doorway was presented. If OPA represents navigational affordances, then stronger responses will be observed when the contralateral visual field is presented with a doorway versus a painting. Bar plots indicate the difference in response to doorways minus paintings. B) Turning direction information (e.g., to the left or right) was also tested based on the visual field (contralateral vs. ipsilateral) of presentation. If OPA represents ego-motion turning, then OPA responses will depend on turn direction, with stronger responses for turns toward the contralateral than ipsilateral visual field. Bar plots indicate the difference in response to contralateral minus ipsilateral turns. C) Forward vs. backward ego-motion information was tested using the whole visual field. If OPA represents ego-motion directions, then different responses will be observed for forward versus backward motion. Bar plots indicate the difference in response to forward minus backward motion. For all plots, error bars indicate the standard error of the mean, and markers indicate data from individual participants.

Open doorways potentially have greater motion energy than paintings, due to changing visual features seen through the doorway as the viewer moves through the scene. To better understand the role of low-level, retinotopic visual properties in driving OPA responses, we next compared responses in OPA to those in EVC. For multivariate analyses, we first tested overall navigational affordance decoding, and found only a marginally significant effect in EVC (t_(15)_ = 1.38, p = 0.09) (Figure 2b). Directly comparing EVC to OPA, a 2 (affordance: within, between) x 2 (region: OPA, EVC) repeated measures ANOVA revealed only a marginally significant affordance x region interaction (F_(1,14)_ = 3.27, p = 0.09). Thus, although we found a trend toward greater sensitivity to affordance information in OPA than EVC, the results of this initial multivariate analysis were inconclusive. We next tested navigational affordance decoding when generalizing across ego-motion directions (i.e., forward vs. backward ego-motion), minimizing the influence of low-level visual features. In this case, navigational affordance decoding in EVC was no longer significant (t_(15)_ = 1.10, p = 0.15), and a 2 (affordance: within, between) x 2 (region: OPA, EVC) repeated measures ANOVA revealed a significant affordance x region interaction (F_(1,14)_ = 4.87, p = 0.045), driven by stronger navigational affordance representation in OPA than EVC. We also performed univariate analyses of responses to contralateral doors vs. paintings. Greater responses to contralateral doors than paintings were not detected in EVC (t_(15)_ = −0.12, p = 0.55), and a 2 (region: OPA, EVC) x 2 (affordance: contralateral, ipsilateral) repeated measures ANOVA revealed a significant region x affordance interaction (F_(1,14)_ = 7.62, p = 0.02), with stronger responses to contralateral affordance information in OPA than EVC (Figure 3a).

The results above provide evidence of a single dissociation in responses between OPA and EVC, with stronger coding of navigational affordances in OPA than EVC. However, stronger evidence that representations in these regions differ would require testing whether the opposite dissociation can also be found; that is, whether there is some information represented more strongly in EVC than OPA. To test this possibility, we considered room texture information. Subjects viewed 8 different room types (counterbalanced across the 10 experimental conditions), which differed based on the textures and colors applied to the walls, floors, and ceiling (i.e., lower-level visual features potentially represented in EVC; Figure 4a). If a region is sensitive to lower-level texture information, then voxelwise patterns of activity will be more similar for stimuli with the same room texture than different room textures. Results from this multivariate analysis showed significant decoding in EVC as well as all three scene regions (EVC: t_(14)_ = 4.24, p < 0.001; OPA: t_(14)_ = 2.89, p = 0.006; PPA: t_(14)_ = 2.34, p = 0.02; MPA: t_(14)_ = 1.93, p = 0.04) (Figure 4b). However, the strength of this effect differed between regions, with significantly stronger sensitivity to room texture information in EVC than OPA (repeated measures ANOVA, region x room type interaction: F_(1,14)_ = 8.26, p = 0.01), as well as PPA (region x room type interaction: F_(1,14)_ = 14.20, p = 0.002) and MPA (region x room type interaction: F_(1,14)_ = 13.92, p = 0.002).

**Figure 4.**
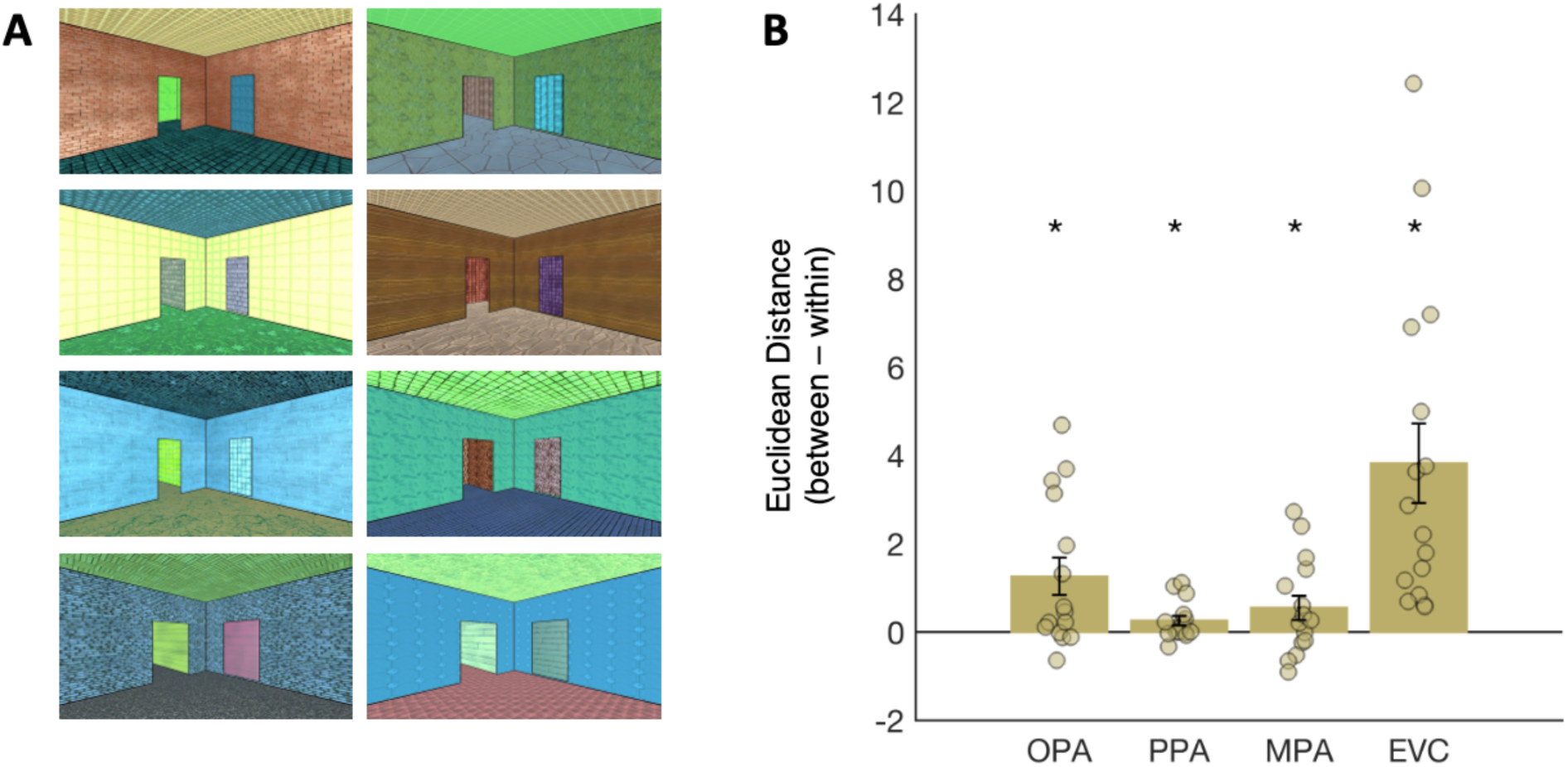
Multivariate analysis of room texture representation. To determine whether OPA responses are dissociable from those in EVC, we tested for information about room textures in each region. Subjects viewed 8 different room types (counterbalanced across conditions), which differed based on the textures and colors applied to the walls, floors, and ceiling. If a region is sensitive to lower-level texture information, then voxelwise patterns of activity will be more similar (Euclidean distance) for stimuli with the same room texture than different room textures. Bar plots indicate the difference score for the same minus the different room type comparisons; responses greater than zero therefore indicate more similar responses (i.e., smaller Euclidean distance) for the “same” than “different” pairs, and thus significant decoding. Error bars represent the standard error of the mean. Markers indicate data from individual participants.

### OPA represents ego-motion in dynamic scenes

If OPA represents ego-motion through dynamic scenes, then OPA responses will differ depending on the direction of ego-motion through the scene. To test this prediction, we analyzed conditions depicting forward (conditions 1-3), backward (conditions (4-6), left turn (conditions 7 and 8), and right turn (conditions 9 and 10) ego-motion. As above, we explored responses to these conditions in a series of multivariate and univariate analyses. For multivariate analyses, we predicted more similar responses (i.e., smaller Euclidean distances) between each of the four ego-motion directions (forward, backward, left, or right) and itself (across split halves of the data) than between different pairs of conditions (also calculated across split halves of the data). Full hypothesis matrices are shown in Figure 2a. Importantly, to minimize visual confounds, between-condition comparisons were limited to the most closely-matched pairs of conditions: left vs. right turn conditions (which were exactly the same stimuli, but mirror flipped) and forward vs. backward conditions (which were exactly the same stimuli, but temporally reversed). Paired samples t-tests revealed significant ego-motion decoding in OPA (t_(14)_ = 3.34, p < 0.001), as well as PPA (t_(14)_ = 2.15, p = 0.003) and MPA (t_(14)_ = 1.28, p = 0.002) (Figure 2b). Directly comparing between regions, a 3 (region: OPA, PPA, MPA) x 2 (ego-motion decoding: within, between) repeated measures ANOVA revealed a significant interaction between region and ego-motion (F_(2, 26)_ = 4.44, p = 0.02), with stronger ego-motion decoding in OPA than PPA (post hoc interaction contrast; F_(1,13)_ = 8.20, p = 0.01), but not MPA (post hoc interaction contrast; F_(1,13)_ = 2.07, p = 0.17). To provide an even stronger test of ego-motion, we also tested ego-motion representation now generalizing across conditions that differed in navigational affordance information. Paired samples t-tests revealed significant ego-motion decoding in OPA (t_(14)_ = 2.01, p = 0.03) and PPA (t_(14)_ = 2.41, p = 0.02), but only a marginally significant trend in MPA (t_(14)_ = 1.57, p = 0.07) (Figure 2b). However, directly comparing between regions, a 3 (region: OPA, PPA, MPA) x 2 (ego-motion decoding: within, between) repeated measures ANOVA failed to reveal a significant interaction between region and ego-motion (F_(2, 26)_ = 2.27, p = 0.12).

To further explore ego-motion representation, we also performed two univariate analyses. Our first analysis tested the prediction that OPA would respond more to forward motion (conditions 1-3) vs. backward motion (conditions 4-6), tested with a paired-samples t-test (two-tailed). Indeed, we found a significantly greater response to forward motion than backward motion in OPA, as well as PPA and MPA (all t’s > 2.52, all p’s < 0.02). Direct comparison between regions revealed a functional dissociation: a 3 (region: OPA, PPA, MPA) x 2 (ego-motion: forward, backward) repeated measures ANOVA revealed a significant region by ego-motion interaction (F_(2, 26)_ = 6.24, p = 0.006), driven by a stronger forward ego-motion preference in OPA than both PPA (F_(1,13)_ = 8.94, p = 0.01) and MPA (F_(1,13)_ = 5.37, p = 0.04).

Our second univariate analysis tested the prediction that responses to ego-motion directions differ by the visual field of presentation, with stronger responses to turns toward the contralateral than ipsilateral visual field (i.e., right OPA responding more to left turns than right turns, and vice versa for left OPA). Note that turns toward the contralateral visual field generate greater motion energy in the ipsilateral field. Accordingly, if a region responds more to contralateral than ipsilateral turns, it is unlikely that this effect is driven by motion energy. To maximize statistical power, responses to contra- and ipsilateral turns were averaged across hemispheres and compared using a paired-samples *t*-test (one-tailed). We found stronger responses to contralateral than ipsilateral turns in OPA (t_(14)_ = 4.14, p < 0.001), as well as PPA (t_(15)_ = 4.22, p < 0.001) and MPA (t_(15)_ = 4.21, p < 0.001) (Figure 3b). Directly comparing between regions, a 3 (region: OPA, PPA, MPA) x 2 (turn: contralateral, ipsilateral) repeated measures ANOVA did not reveal a significant interaction between region and ego-motion (F_(2, 26)_= 2.53, p = 0.098), despite trends in the predicted direction.

We next compared ego-motion representation in OPA with that in EVC. For multivariate analyses, we first tested overall ego-motion decoding, and found significant sensitivity to ego-motion direction in EVC (t_(14)_ = 1.28, p = 0.001), with no significant difference in the strength of ego-motion decoding between OPA and EVC; a 2 (region: OPA, EVC) x 2 (ego-motion decoding: within, between) repeated measures ANOVA failed to find a significant ego-motion by region interaction (F_(1,14)_ = 0.07, p = 0.80) (Figure 2b). Next, we tested ego-motion direction information generalizing across navigational affordance information, providing a stronger test of ego-motion representation over and above low-level visual features. In this analysis, decoding of ego-motion direction was no longer significant in EVC (t_(14)_ = 0.05, p = 0.52), although direct comparison of OPA and EVC using a 2 (region: OPA, EVC) x 2 (ego-motion decoding: within, between) repeated measures ANOVA revealed only a marginally significant region by ego-motion interaction (F_(1,14)_ = 3.24, p = 0.09), with a trend toward stronger coding in OPA than EVC (Figure 2b). Finally, we performed two univariate analyses. First, for univariate analyses of forward vs. backward motion responses, we found a preference for forward over backward motion in EVC (paired samples t-test, t(15) = 2.89, p = 0.01), with no difference in response between OPA and EVC (region by ego-motion interaction; F(1,14) = 0.38, p = 0.55) (Figure 3c). Second, for univariate analyses comparing contralateral vs. ipsilateral turns, we found no greater response to contra-than ipsilaterally-presented turns in EVC (t_(15)_ = 0.39, p = 0.90), and a 2 (region: OPA, EVC) x 2 (turn: contralateral, ipsilateral) repeated measures ANOVA revealed a significant region x turn interaction (F_(1,14)_ = 15.65, p = 0.001) (Figure 3b). Taken together, these results suggest that OPA representation of ego-motion may be partially, but not entirely explained by low-level, retinotopic visual information. Indeed, although EVC responses can discriminate between basic ego-motion directions, the full profile of OPA responses to ego-motion is not detectable in EVC – particularly for tests of ego-motion representation that minimize the influence of low-level visual features (e.g., when generalizing across different navigational affordance conditions, or for contralateral vs. ipsilateral turns).

### Integration of navigational affordances and ego-motion

The analyses above suggest that OPA plays a unique role in the scene network by representing both navigational affordances and ego-motion in dynamic scenes. Does OPA integrate these two kinds of navigationally relevant information? To begin to address this question, we compared responses to conditions in which navigational affordance and ego-motion information are consistent (e.g., an open doorway to the left, with a turn toward the door; conditions 7 and 10) versus inconsistent (e.g., an open doorway to the left, with a turn away from the door; conditions 8 and 9) (Figure 5a). Once again, we tested responses to these conditions with a series of multivariate and univariate analyses.

**Figure 5.**
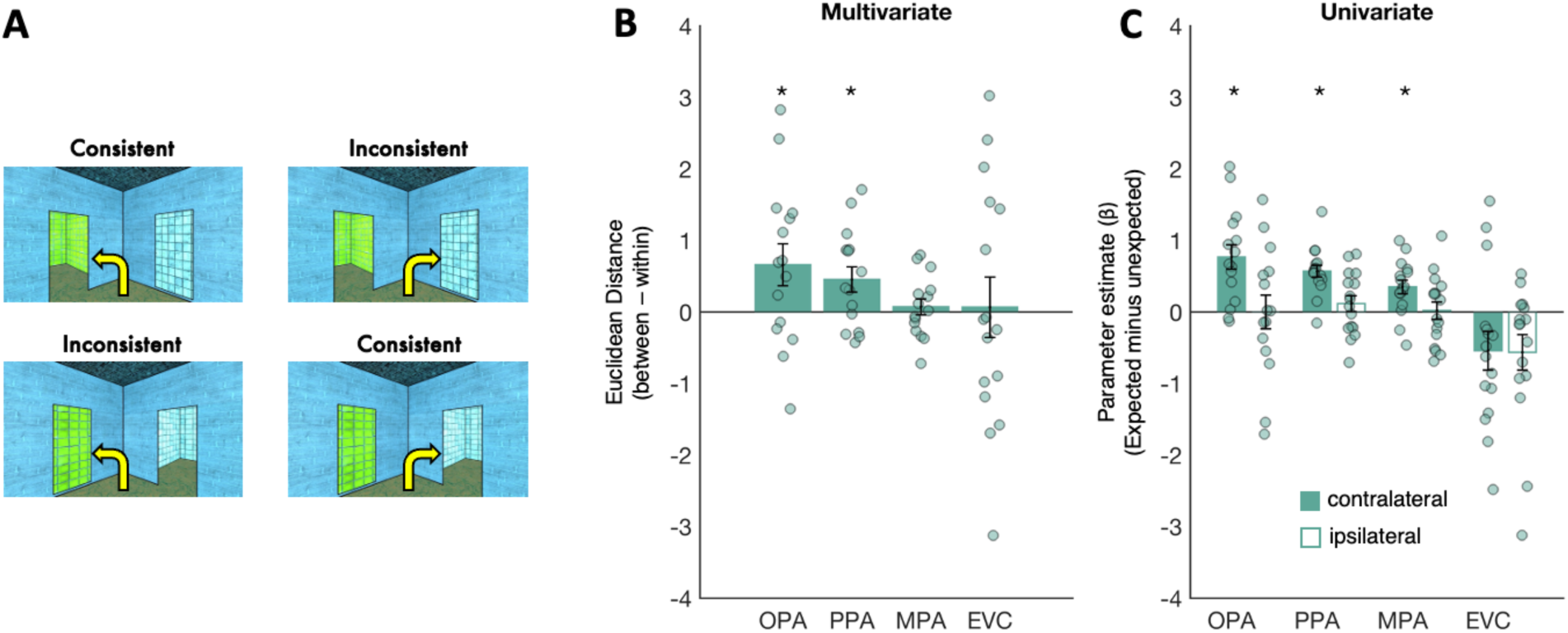
Testing the conflict of navigational affordance and ego-motion information. A) Conditions used for conflict analyses. Navigational affordance and ego-motion are consistent when turns are taken in the direction of the open doorway (top left and bottom right conditions). This information is put in conflict when turns are taken in the opposite direction of the doorway (top right and bottom left conditions). B) Multivariate analyses. If OPA represents the conflict of navigational affordance and ego-motion information, then more similar responses will be observed for conditions with the same conflict information (consistent-consistent, inconsistent-inconsistent) than different conflict information (consistent-inconsistent). Bar plots indicate the difference score for the “between” conditions minus the “within” conditions; responses greater than zero therefore indicate more similar responses (i.e., smaller Euclidean distance) for the “within” than “between” condition pairs, and thus significant decoding. C) Univariate analysis were conducted with respect to the visual field of presentation. If OPA represents the conflict of navigational affordance and ego-motion information, specifically in the contralateral visual field, then stronger responses to consistent than inconsistent turns will be found for contralateral turns, but not ipsilateral turns. Bar plots indicate the difference in response to consistent minus inconsistent turns. For all plots, error bars represent the standard error of the mean. Markers indicate data from individual participants.

For multivariate analyses, we predicted more similar responses within consistent and inconsistent conditions (across split halves of the data) than between these conditions (also calculated across split halves of the data) (Figure 5b). Paired samples t-tests revealed a significant effect in OPA (t_(14)_ = 2.26, p = 0.02) and PPA (t_(14)_ = 2.56, p = 0.01), but not MPA (t_(14)_= 0.65, p = 0.26). A 3 (region: OPA, PPA, MPA) x 2 (decoding: within, between) repeated measures ANOVA did not find a significant region x decoding interaction (F_(1.26, 16.35)_ = 2.07, p = 0.17). Notably, the differential response to consistent versus inconsistent turning conditions was not likely explained by low-level, retinotopic visual features, since no significant effect was found in EVC (paired samples t-test, t_(15)_ = 0.16, p = 0.44), although direct comparison between EVC and OPA did not reveal a significant region x decoding interaction (F_(1,14)_ = 0.92, p = 0.35).

To further explore how navigational affordance and ego-motion information are integrated, we next tested the univariate prediction that OPA would respond differently to consistent versus inconsistent turns, particularly when this information is presented in the contralateral visual field (e.g., for right OPA, a preference for left turns toward an open doorway versus left turns toward a painting). Consistent with this prediction, we found greater OPA responses to consistent than inconsistent turns toward the contralateral visual field (paired samples t-test, t_(14)_ = 4.50, p < 0.0001). This effect was not likely explained by a simple preference for doors versus paintings in the contralateral visual field, since no difference in response was found for consistent vs. inconsistent ipsilateral turns (paired samples t-test, t_(14)_ = −0.001, p = 0.50); if OPA is simply responding to doorways in the contralateral visual field, then OPA would respond significantly more to inconsistent, ipsilateral turns (which contain a doorway in the contralateral visual field) than consistent, ipsilateral turns (which contain a painting in the contralateral visual field). The preference for consistent contralateral turns was also observed in PPA and MPA (paired samples t-tests; contralateral: both t’s_(15)_ > 3.68, both p’s < 0.002; ipsilateral: both t’s_(15)_ = 1.06, p = 0.15). To directly compare between regions, we first calculated the difference in response to the consistent minus the inconsistent condition in each visual field, and then submitted these values to a 2 (visual field: contralateral, ipsilateral) x 3 (region: OPA, PPA, MPA) repeated measures ANOVA. This analysis found only a marginally significant interaction between visual field and region (F_(2, 28)_ = 2.53, p = 0.098), with no difference in preference for consistent, contralateral turns in OPA vs. PPA (interaction contrast, F_(1,14)_ = 1.62, p = 0.22), and a marginally significant trend toward a stronger preference in OPA than MPA (interaction contrast, F_(1,14)_ = 3.90, p = 0.07). In contrast to the scene regions, EVC did not respond more consistent versus inconsistent turns in either the contralateral (paired samples t-test, t_(15)_ = −2.01, p = 0.97) or the ipsilateral visual field (paired samples t-test, t_(15)_ = −2.26, p = 0.98). In sum, the scene regions, and particularly OPA, integrate navigational affordance and ego-motion information, and respond maximally when they converge.

## Discussion

Here we used dynamic movie stimuli to test how navigationally relevant information is represented in scene-selective cortex during the first-person visual experience of navigating scenes. We found that one scene region, the OPA, was sensitive to both navigational affordances (i.e., the presence versus absence of open doorways affording further navigation) and ego-motion directions (i.e., walking forward vs. backward, turning left vs. right). Weaker and less consistent results were observed in two other scene regions, the PPA and MPA, which showed no sensitivity to navigational affordance information, and relatively weaker evidence of ego-motion representation. These results support the hypothesis that regions of the cortical scene processing system are functionally dissociated, with OPA playing a specific role in representing navigationally relevant information in dynamic visual scenes.

Our results reveal a more detailed functional profile for OPA than previously known. While past work had shown that OPA is activated more strongly by dynamic than static scenes (Hacialihafiz and Bartels, 2015; Kamps et al., 2016b; Kamps et al., 2020; Sulpizio et al., 2020; Suzuki, Kamps et al., 2021), here we explored the more specific representations in OPA during the visual experience of navigating through a scene. In so doing, we found that OPA represents information about both the structure of the space as it constrains possibilities for future navigation (i.e., navigational affordances), the dynamics of ongoing motion through the space (i.e., ego-motion), and their interaction. Given that navigational affordance information is particularly useful for understanding possibilities for future actions (e.g., which way can I walk next?) while ego-motion reveals the current trajectory of motion (e.g., how am I currently moving?), these results suggest that OPA is involved not only in navigational planning, but also in online guidance of navigation.

Prior work has typically studied (static) scene perception and ego-motion perception separately, with studies of static scene perception focused on the three regions tested here (OPA, PPA, and MPA), and studies of ego-motion focused on a network of regions in dorsal visual cortex, parietal cortex, premotor cortex, and the cerebellum (e.g., Wall and Smith, 2008; Sulpizio et al., 2020; Di Marco, et al., 2021; Ruehl et al., 2022; Pitzalis et al., 2020; Sulpizio et al., 2023). The current results support the idea that OPA may play a relatively unique role at the intersection of these two networks (Sulpizio et al., 2020), by representing information about navigational affordances – a property of static scene geometry – and ego-motion direction - a property only defined in dynamic scenes. We speculate that the co-localization of these two types of information in OPA might support navigation by facilitating comparison between planned navigational trajectories and currently observed progress in following that trajectory. Supporting this idea, we found initial evidence that OPA is sensitive to the conflict of navigational affordance and ego-motion information (e.g., turning toward versus away from an open doorway) – although this effect was not specific to OPA, but also found in other scene regions.

Although OPA is sensitive to higher-level information relevant to navigation, OPA responses are nevertheless constrained by the visual field of presentation. Mirroring the broader organization of early visual processing, past studies have demonstrated a clear preference in OPA for contralateral visual stimulation, particularly in the upper visual field (MacEvoy and Epstein, 2007; Silson et al., 2015; Silson et al., 2022). Silson et al. (2015) further showed that visual field biases in OPA (and other scene regions) are stronger when mapped with fragments of static scene images, compared with simple checkerboard stimuli, suggesting that OPA encodes high-level representations of scene stimuli, particularly in the upper, contralateral visual field. Our results build on this finding by revealing the more specific scene information extracted by OPA in this portion of the visual field, namely, navigational affordances and ego-motion. Future work will be required to explore how the navigationally relevant information encoded separately in each hemisphere is integrated to form a coherent representation across the entire visual field, for example, facilitating decision-making about which paths to follow to the left vs. right.

While ego-motion representation was found in OPA, at least some sensitivity to this information was found in all other regions tested, including EVC. Both OPA and EVC further showed a preference for forward (vs. backward) ego-motion, suggesting that ego-motion representation is biased toward the direction most commonly experienced during visually-guided navigation. Widespread representation of ego-motion across the visual cortex is perhaps not surprising, given that ego-motion directions generate distinct patterns of optic flow across the visual field (e.g., a left turn causes higher motion energy in the right visual field, and vice versa for a right turn). Nevertheless, some evidence indicates that nature of ego-motion representation differs between OPA and EVC. For example, ego-motion decoding remained significant in OPA, but not EVC, when generalizing across stimuli that differed in navigational affordance information (a test which minimizes the influence of lower-level motion energy in driving decoding performance). Moreover, univariate responses in OPA were greater for turns in the contralateral direction (e.g., left OPA responding more to right turns than left turns). If responses in OPA were driven only by lower-level motion energy, then greater responses should be found for ipsilateral turns, which generate greater motion energy in contralateral visual field. Indeed, and by contrast, EVC responded more to ipsilateral than contralateral turns. Based on these results, we speculate that OPA encodes relatively higher-level representations of ego-motion, perhaps reflecting ongoing navigational goals, while ego-motion representation in EVC is driven more directly by low-level motion energy information.

In sum, we used dynamic visual stimuli to find evidence of a functional dissociation in the human cortical scene network, with OPA showing stronger and more consistent evidence of navigationally-relevant information processing in dynamic scenes than PPA or MPA. Moreover, the pattern of responses in OPA could not be explained by low-level, retinotopic visual features. These results support the hypothesis that OPA plays a unique role in the cortical scene processing network by representing scenes as dynamic, navigable spaces.

## Methods

### Participants

Sixteen participants (Mean age = 29.3 years, range = 20–45 years; 9 females) were recruited from the MIT community. All participants gave written informed consent to participate and had normal or corrected-to-normal vision. The sample size was preregistered prior to analysis, and was selected on the basis of prior fMRI studies testing representation of navigational affordances in OPA (Bonner and Epstein, 2017; Park and Park, 2020). A power analysis for the critical paired-samples t-tests suggested that a sample size of 16 provided 85% power to detect large effects (i.e., d = 0.8; if we assumed the even larger effect sizes reported in previous work (e.g., d = 1.02 in Bonner et al., 2017), power increased to 97%). Recruitment and experiment protocols were approved by the Committee on the Use of Humans as Experimental Subjects (COUHES) at the Massachusetts Institute of Technology.

### Design

We used a region of interest (ROI) approach in which we used one set of runs (“Localizer Runs”) to localize ROIs based on functional signatures, and an distinct independent set of runs (“Experimental Runs”) to investigate responses of these regions to the Experimental conditions shown in Figure 1, using both univariate and multivariate approaches (for a detailed description of this analysis see Data analysis section below).

Localizer stimuli consisted of 3s videos of dynamic Scenes, Objects, Faces, and Scrambled Objects, as described previously in Kamps et al., 2016 and 2020. Stimuli were presented using a block design at 13.7 x 18.1 degrees of visual angle. Each run was 315s long and contained 4 blocks per stimulus category. The order of the first set of blocks was pseudorandomized across runs (e.g., Faces, Objects, Scenes, Scrambled) and the order of the second set of blocks was the palindrome of the first (e.g., Scrambled, Scenes, Objects, Faces). Each block consisted of 5 2.8s video clips from a single condition, with an ISI of 0.2s, resulting in 15s blocks. Each run also included 5 fixation blocks: one at the beginning, three evenly spaced throughout the run, and one at the end. Participants completed 3 Localizer runs, interleaved between every 2 Experimental Runs.

Experimental stimuli consisted of 14 conditions (Conditions 1-10 are shown in Figure 1; Example stimuli and results from conditions 11-14 are shown in Supplemental Figure 2). All stimuli were created using Unity software and depicted 3 second clips of the first-person experience of walking through scenes. Navigational affordances were manipulated by including an open doorway to either the left side, right side, or both sides. To help control for low-level visual confounds, the non-doorway side always included a distractor object, either a painting (conditions 1-10) or an inverted doorway (conditions 11-14). Furthermore, the textures applied to the painting and the walls through the doorways were counterbalanced, such that each texture appeared equally on either side across the full stimulus set. Ego-motion was manipulated by changing the direction of ego-motion through scene, which could either be forward (conditions 1-3, 11-14), backward (conditions 4-6), a left turn (conditions 7-8) or a right turn (conditions 9-10). To help prevent visual adaptation over the course of the experiment, the 14 experimental conditions were counterbalanced across 8 room types, which differed from one another based on the textures applied to the walls, floor and ceiling, and to a lesser extent, by the size and shape of the doorways and corresponding distractor (Figure 4). Stimuli were presented at 13.1 x 18.6 DVA in an event-related paradigm. Each stimulus was presented for 2.5s, followed by a minimum inter-stimulus-interval (ISI) of 3.5s and a maximum ISI of 9.5s, optimized separately for each run using OptSeq2. Participants viewed 4 repetitions of each condition per run, and completed 8 experimental runs, yielding 32 total repetitions per condition across the experiment. Participants performed a one-back task to help ensure attention throughout the experiment, and were instructed to lie still, keep their eyes open, and try to “pay attention to” and “immerse themselves in” the stimuli.

### Data acquisition

Data were acquired from a 3-Tesla Siemens Magnetom Prisma scanner located at the Athinoula A. Martinos Imaging Center at MIT, using a 32-channel head coil. Participants viewed movie stimuli through a mirror projected to a screen behind the scanner. Scout images (3D low-resolution anatomical scans) were acquired using auto-align in 128 sagittal slices with 1.6mm isotropic voxels (TA=0.14; TR=3.15ms; FOV=260mm). Anatomical T1-weighted structural images were acquired in 176 interleaved sagittal slices with 1.0mm isotropic voxels (MPRAGE; TA=5:53; TR=2530.0ms; FOV=256mm; GRAPPA parallel imaging, acceleration factor of 2). Functional data were acquired with a gradient-echo EPI sequence sensitive to Blood Oxygenation Level Dependent (BOLD) contrast in 2mm isotropic voxels in 46 interleaved near-axial slices covering the whole brain (EPI factor=70; TR=2s; TE=30.0ms; flip angle=90 degrees; FOV=210mm). For the Localizer runs, 158 volumes were acquired per run (TA=5:16). For the Experimental runs, 228 volumes were acquired per run (TA=7:36). Due to experimenter and technical errors, the actual number of volumes collected occasionally deviated from these values. In these cases, all timepoints were included in the analysis, with only usable trials included in the GLM.

### Preprocessing

Results included in this manuscript come from preprocessing performed using fMRIPrep 21.0.1 (Esteban, Markiewicz, et al. (2018); Esteban, Blair, et al. (2018); RRID:SCR_016216), which is based on Nipype 1.6.1 (K. Gorgolewski et al. (2011); K. J. Gorgolewski et al. (2018); RRID:SCR_002502).

#### Anatomical data preprocessing

One T1-weighted (T1w) image was collected per participant. The T1-weighted (T1w) image was corrected for intensity non-uniformity (INU) with N4BiasFieldCorrection (Tustison et al. 2010), distributed with ANTs 2.3.3 (Avants et al. 2008, RRID:SCR_004757), and used as T1w-reference throughout the workflow. The T1w-reference was then skull-stripped with a Nipype implementation of the antsBrainExtraction.sh workflow (from ANTs), using OASIS30ANTs as target template. Brain tissue segmentation of cerebrospinal fluid (CSF), white-matter (WM) and gray-matter (GM) was performed on the brain-extracted T1w using fast (FSL 6.0.5.1:57b01774, RRID:SCR_002823, Zhang, Brady, and Smith 2001). Brain surfaces were reconstructed using recon-all (FreeSurfer 6.0.1, RRID:SCR_001847, Dale, Fischl, and Sereno 1999), and the brain mask estimated previously was refined with a custom variation of the method to reconcile ANTs-derived and FreeSurfer-derived segmentations of the cortical gray-matter of Mindboggle (RRID:SCR_002438, Klein et al. 2017). Volume-based spatial normalization to two standard spaces (MNI152NLin6Asym, MNI152NLin2009cAsym) was performed through nonlinear registration with antsRegistration (ANTs 2.3.3), using brain-extracted versions of both T1w reference and the T1w template. The following templates were selected for spatial normalization: FSL\u2019s MNI ICBM 152 non-linear 6th Generation Asymmetric Average Brain Stereotaxic Registration Model [Evans et al. (2012), RRID:SCR_002823; TemplateFlow ID: MNI152NLin6Asym], ICBM 152 Nonlinear Asymmetrical template version 2009c [Fonov et al. (2009), RRID:SCR_008796; TemplateFlow ID: MNI152NLin2009cAsym].

#### Functional data preprocessing

For each of the 9-12 BOLD runs per subject (across all tasks and sessions), the following preprocessing was performed. First, a reference volume and its skull-stripped version were generated using a custom methodology of fMRIPrep. Head-motion parameters with respect to the BOLD reference (transformation matrices, and six corresponding rotation and translation parameters) are estimated before any spatiotemporal filtering using mcflirt (FSL 6.0.5.1:57b01774, Jenkinson et al. 2002). The BOLD time-series (including slice-timing correction when applied) were resampled onto their original, native space by applying the transforms to correct for head-motion. These resampled BOLD time-series will be referred to as preprocessed BOLD in original space, or just preprocessed BOLD. The BOLD reference was then co-registered to the T1w reference using bbregister (FreeSurfer) which implements boundary-based registration (Greve and Fischl 2009). Co-registration was configured with six degrees of freedom. Several confounding time-series were calculated based on the preprocessed BOLD: framewise displacement (FD), DVARS and three region-wise global signals. FD was computed using two formulations following Power (absolute sum of relative motions, Power et al. (2014)) and Jenkinson (relative root mean square displacement between affines, Jenkinson et al. (2002)). FD and DVARS are calculated for each functional run, both using their implementations in Nipype (following the definitions by Power et al. 2014). The three global signals are extracted within the CSF, the WM, and the whole-brain masks. Additionally, a set of physiological regressors were extracted to allow for component-based noise correction (CompCor, Behzadi et al. 2007). Principal components are estimated after high-pass filtering the preprocessed BOLD time-series (using a discrete cosine filter with 128s cut-off) for the two CompCor variants: temporal (tCompCor) and anatomical (aCompCor). tCompCor components are then calculated from the top 2% variable voxels within the brain mask. For aCompCor, three probabilistic masks (CSF, WM and combined CSF+WM) are generated in anatomical space. The implementation differs from that of Behzadi et al. in that instead of eroding the masks by 2 pixels on BOLD space, the aCompCor masks are subtracted a mask of pixels that likely contain a volume fraction of GM. This mask is obtained by dilating a GM mask extracted from the FreeSurfer (2019) aseg segmentation, and it ensures components are not extracted from voxels containing a minimal fraction of GM. Finally, these masks are resampled into BOLD space and binarized by thresholding at 0.99 (as in the original implementation). Components are also calculated separately within the WM and CSF masks. For each CompCor decomposition, the k components with the largest singular values are retained, such that the retained components\u2019 time series are sufficient to explain 50 percent of variance across the nuisance mask (CSF, WM, combined, or temporal). The remaining components are dropped from consideration. The head-motion estimates calculated in the correction step were also placed within the corresponding confounds file. The confound time series derived from head motion estimates and global signals were expanded with the inclusion of temporal derivatives and quadratic terms for each (Satterthwaite et al. 2013). Frames that exceeded a threshold of 0.5 mm FD or 1.5 standardised DVARS were annotated as motion outliers. The BOLD time-series were resampled into standard space, generating a preprocessed BOLD run in MNI152NLin6Asym space. First, a reference volume and its skull-stripped version were generated using a custom methodology of fMRIPrep. Automatic removal of motion artifacts using independent component analysis (ICA-AROMA, Pruim et al. 2015) was performed on the preprocessed BOLD on MNI space time-series after removal of non-steady state volumes and spatial smoothing with an isotropic, Gaussian kernel of 6mm FWHM (full-width half-maximum). Corresponding non-aggresively denoised runs were produced after such smoothing. Additionally, the aggressive noise-regressors were collected and placed in the corresponding confounds file. All resamplings can be performed with a single interpolation step by composing all the pertinent transformations (i.e. head-motion transform matrices, susceptibility distortion correction when available, and co-registrations to anatomical and output spaces). Gridded (volumetric) resamplings were performed using antsApplyTransforms (ANTs), configured with Lanczos interpolation to minimize the smoothing effects of other kernels (Lanczos 1964). Non-gridded (surface) resamplings were performed using mri_vol2surf (FreeSurfer).

Many internal operations of fMRIPrep use Nilearn 0.8.1 (Abraham et al. 2014, RRID:SCR_001362), mostly within the functional processing workflow. For more details of the pipeline, see the section corresponding to workflows in fMRIPrep (2019) documentation. The above boilerplate text was automatically generated by fMRIPrep with the express intention that users should copy and paste this text into their manuscripts unchanged. It is released under the CC0 license.

### Modeling

For run-level analyses, the preprocessed timeseries were assessed with algorithms from the Artifact Removal Toolbox (ART) to identify frames within the run that had an abnormal amount of motion (0.4 mm of total displacement, or an intensity spike >3 s.d. from the mean). The design matrix included boxcars for the experimental conditions convolved with a double-gamma hemodynamic response function (HRF), and nuisance regressors representing framewise motion, the anatomical CompCor regressors derived from white matter and cerebrospinal fluid, as well as impulse regressors for volumes identified by ART. A high-pass filter (120 Hz) was applied to the design matrix and the smoothed data. The model was evaluated using FSL’s FILM program. Subject-level contrast maps will be generated using FSL’s FLAME in fixed-effects mode.

### Data exclusion

For one participant, two runs were lost due to an experimenter error that prevented recovery of event timings, resulting in 6 usable runs. For three participants, 3-5 runs were excluded due to excessive motion. For five participants, there was one run in which the script froze toward the end of the run. These runs were kept, and events were modeled up to the point at which the script froze. Following these exclusions, participants had either three (N=14), two (N=1), or one (N=1) usable localizer run, and either eight (N=12), six (N=1), or five (N=3) usable experimental runs.

### ROI definition

ROIs included three scene-selective regions: the occipital place area (OPA), parahippocampal place area (PPA), and medial place area (MPA), as well as one early visual cortex control region (EVC). We first hand-defined a contiguous cluster of scene-selective voxels for each scene region in each subject based on the contrast of scenes > objects. The localizer data were thresholded at p < 10^-3^, and only voxels surviving this threshold were included. OPA, PPA, and MPA were then defined as the top 50 voxels within the hand-defined mask in each hemisphere. All regions were defined in all participants, except for one subject missing OPA, and a second subject missing both PPA and RSC. For EVC, we used a group-constrained, subject-specific (GSS) method (Julian et al., 2012) in which we selected the top 50 voxels from each hemisphere within a larger anatomical search space taken from Wang et al. (2015). Top EVC voxels were ranked and selected based on the contrast of all conditions > fixation.

### Univariate analyses

For univariate analyses, responses of all voxels in each ROI were averaged together for each run, and then averaged across runs, yielding the overall response of each ROI in each participant to each experimental condition. Responses were extracted separately from each hemisphere, and then averaged across hemispheres for all analyses, except where hemispheric differences were expected based on the visual field (contralateral versus ipsilateral) of presentation (see Results).

### Multivariate analyses

For multivariate analysis, we used a split-half procedure in which the average response from each voxel was calculated across all possible combinations of half of the usable runs (thus leaving out a complementary, independent half of the runs for each fold of the data). For each fold, we estimated the similarity of voxel-wise patterns of activity between conditions by calculating the Euclidean distance between each condition and every other condition (including itself) in the other half of the data. These estimates were averaged across all folds, yielding an overall estimate of the condition-wise similarity space for each ROI and participant.

## Data and code availability

The methods and analyses of this study were pre-registered prior to data analysis. Deviations from the pre-registered analyses, and results from originally planned analyses, are detailed in the supplemental materials. Preregistration documents, experiment scripts, stimuli, data, and analysis scripts required to produce statistical results are available at https://osf.io/6yehp/?view_only=53ba7725d61343a29a2e3e6da5d75f28, and will be made publicly available prior to publication. De-faced brain images from participants who consented to share them will also be made available on OpenNeuro by the time of publication.

## Author contributions

F.K., R.S., and N.K. designed the research. F.K. and E.C. collected and analyzed the data. F.K. wrote the paper with feedback from E.C., N.K., and R.S.

## Acknowledgements

We thank: Atsushi Takahashi, Steve Shannon, and the Athinoula A. Martinos Imaging Center at the McGovern Institute at MIT for technical and administrative support; Kirsten Lydic for technical support; and members of the Saxe and Kanwisher labs for feedback and assistance with data collection.

## Supplemental materials

### Deviations from preregistered analyses

Our primary analyses (as reported in the manuscript) deviated from our preregistered analyses in three ways. First, many of our preregistered analyses depended on comparisons between single pairs of conditions. After seeing the results of the preregistered analyses, we decided to perform the analyses reported in the main results of the manuscript, which involved averaging responses across a greater number of conditions, in order to increase power to detect key effects. Second, we initially planned to define scene ROIs using a group-constrained, subject specific (GSS) method, but after visually inspecting the localizer data, we discovered that the peak OPA activation often fell outside the parcel search space. To ensure accurate ROI definition, we therefore opted to define the scene ROIs by hand, as described in the manuscript. Third, and finally, we analyzed data expressed as raw betas, rather than percent signal change. For transparency, results of all original, pre-registered analyses not reported in the main results are presented below. Notably, we still opted to report these analyses in hand-defined ROIs, rather than GSS ROIs, since the GSS ROIs were determined to be inaccurate. Results are also reported as raw betas to facilitate comparison to the primary analyses. In our view, these deviations from the preregistered analysis plan are minor and well-justified. Nevertheless, because we made these changes after conducting some initial analyses, the results reported here must be considered exploratory, and should be replicated in a fully independent sample.

### Additional preregistered analyses of navigational affordance representation

If OPA represents navigational affordances, then OPA responses will differ depending on the number and direction of navigational affordances present in the scene. To test this prediction, we analyzed conditions 1, 2, and 3 only, which depict forward motion through scenes with either i) an open doorway to the left (condition 2); ii) an open doorway to the right (condition 3); or iii) two open doorways (condition 1). For univariate analyses, we found no evidence of differential responses to 2-door versus 1-door scenes (calculated as the average of left-and right-door scenes; two-tailed paired samples t-test) in OPA (t_(14)_ = 1.35, p = 0.20), nor in PPA (t_(14)_ = 1.86, p = 0.08), MPA (t_(14)_ = 0.30, p = 0.77), or EVC (t_(15)_ = 1.57, p = 0.14) (Supplemental Figure 1). For multivariate analyses, we predicted stronger correlations between each of the three affordance conditions (door to the left, right, or both) and itself (across split halves of the data) than between each condition and each other condition (also calculated across split halves of the data), tested with a paired samples *t*-test (one-tailed). We found a marginally significant effect in OPA (t_(14)_ = 1.39, p = 0.09) and PPA (t_(14)_ = 1.55, p = 0.07), and no significant difference in MPA (t_(14)_ = 0.80, p = 0.22) or EVC (t_(15)_ = 1.25, p = 0.12) (Supplemental Figure 1). Direct comparison between regions failed to find a difference between OPA and all other regions (tested with repeated measures ANOVA; all F’s < 0.62, all p’s > 0.46).

**Supplemental Figure 1.**
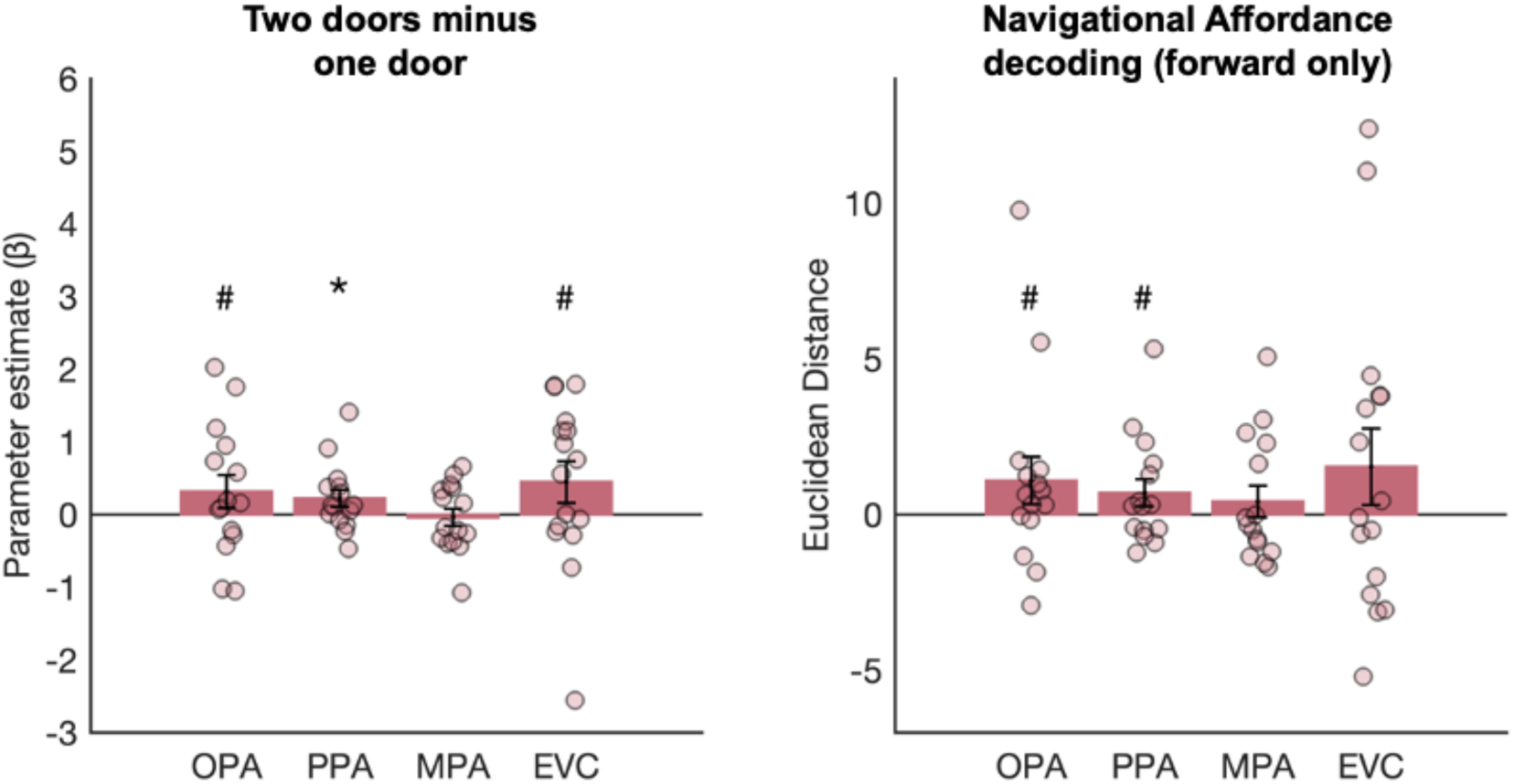
Additional tests of navigational affordance representation. A) Univariate responses (difference scores) to two door minus one door scenes, tested using only the forward motion conditions (i.e., conditions 1-3). B) Multivariate decoding of navigational affordances tested only using forward motion conditions (i.e., conditions 1-3). For both plots, error bars represent the standard error of the mean. Markers indicate data from individual participants.

Positive findings in the analyses above could have been driven by a confound of lower-level dynamic visual information, since ego-motion through space will cause greater changes in the dynamic patterns of occlusion visible through the doorway, as compared with the painting. To help address this possibility, we analyzed conditions 11 and 12, which are similar to conditions 2 and 3, but depict an inverted doorway on the control side (rather than a painting) (Supplemental Figure 2). The inverted doorway is precisely matched to the open doorway in terms of lower-level dynamic visual information, except that it cannot be navigated via typical, bipedal locomotion. We made two multivariate predictions. First, in order to demonstrate basic sensitivity to the direction of the navigational affordance in these conditions, we predicted that patterns of activation would be more similar between each condition and itself (across split halves of the data) than between each condition and the other condition, tested with a paired samples *t*-test (one-tailed). We found a marginally significant effect in OPA (t_(14)_ = 1.42, p = 0.09), significant effects in PPA (t_(14)_ = 1.93, p = 0.04) and EVC (t_(15)_ = 3.10, p = 0.004), and no significant effect in MPA (t_(14)_ = 0.66, p = 0.26). The effect in OPA was significantly weaker than that in EVC (F_(1,14)_ = 5.38, p = 0.04), and did not significantly differ from that in PPA or MPA (both Fs < 2.82; both p’s > 0.11). Second, we predicted that if OPA represents navigational affordances – not just low-level dynamic visual information – then coding of navigational affordance information will generalize across scene features that place similar constraints on navigation, but which differ in terms of dynamic visual information (i.e., paintings versus inverted doors). Specifically, we predicted stronger correlations across conditions 2-11 and 3-12 than across conditions 2-12 and 3-11, tested with a paired samples *t*-test (one-tailed). This analysis failed to reveal a significant effect in OPA (t_(14)_ = 0.10, p = 0.54), MPA (t_(14)_ = 0.40, p = 0.65), and EVC (t_(15)_ = 0.09, p = 0.46), although a significant effect was observed in PPA (t_(14)_ = 1.96, p = 0.03). However, no significant differences were found between the three scene regions (repeated measures ANOVA, region x condition interaction: F_(2,26)_ = 1.36, p = 0.28) or between OPA and EVC (region x condition interaction: F_(1,14)_ = 0.05, p = 0.83). Finally, to further evaluate the robustness of navigational affordance coding in OPA, we included two further inverted door conditions (conditions 13 and 14), which depicted forward motion through rooms with a different overall spatial layout than that in the conditions above (Supplemental Figure 2). If OPA represents navigational affordances, then patterns of activity should be more similar between each condition and itself (across split halves of the data) than between each condition and the other condition, tested with a paired samples *t*-test (one-tailed). Again, we failed to find a significant effect in OPA (t_(14)_ = 0.19, p = 0.43), although a significant effect was detected in PPA (t_(14)_ = 2.01, p = 0.03), and marginally significant effects were detected in MPA (t_(14)_ = 1.36, p = 0.098) and EVC (t_(14)_ = 01.57, p = 0.07).

**Supplemental Figure 2.**
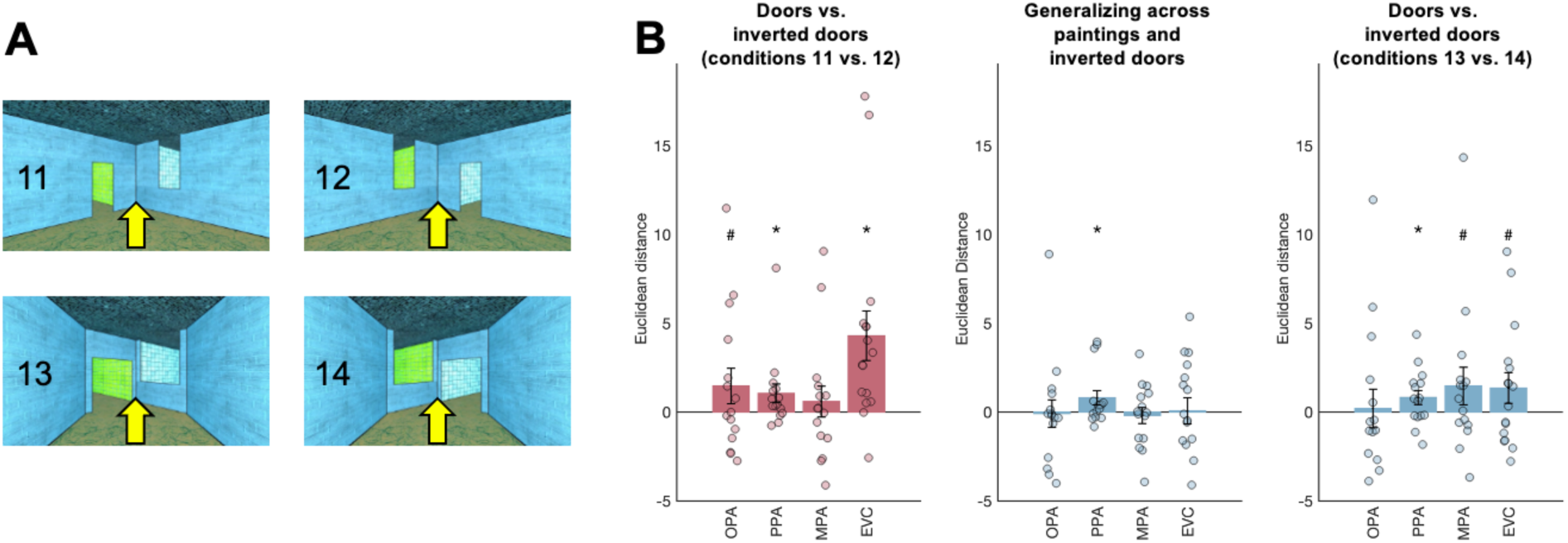
Testing responses to dooways vs. inverted doorways. A) Example stimuli from conditions 11-14. B) Multivariate decoding results from the inverted doorway conditions. Bar plots indicate the difference score for the “between” conditions minus the “within” conditions; responses greater than zero indicate more similar responses (i.e., smaller Euclidean distance) for the “within” than “between” condition pairs, and thus significant decoding. Error bars represent the standard error of the mean. Markers indicate data from individual participants.

### Additional preregistered analyses of ego-motion representation

If OPA represents navigational dynamics, then OPA responses will differ depending on the direction of ego-motion through the scene. Key pre-registered analyses testing this hypothesis are already presented in the main results. For an additional multivariate analyses, we analyzed forward (conditions 1-3) and backward (conditions 4-6) motion conditions, and predicted stronger correlations across conditions that share a direction of motion (forward-forward or backward-backward, across split halves of the data, and across all affordances) than conditions that differ in the direction of motion (i.e., forward-backward) (Supplemental Figure 3). A paired samples t-test (one-tailed) revealed a significant difference in OPA (t_(14)_ = 2.02, p = 0.03) but not PPA, MPA, or EVC (all t’s < 1.63, all p’s > 0.06). However, repeated measures ANOVAs with factors for region (OPA, PPA, and MPA; or OPA and EVC) and condition (within, between) failed to reveal significant region x condition interactions (both F’s < 1.85, both p’s > 0.17).

Second, we pre-registered a multivariate analysis comparing responses to scene movies depicting left turns (conditions 2 and 3) versus right turns (conditions 5 and 6). We predicted stronger correlations across conditions depicting the same direction of motion (left-left, right-right, across split halves of the data) than those that differ in the direction of motion (left-right). Paired samples t-tests (one-tailed) revealed a marginally significant difference in OPA (t_(14)_ = 1.62, p = 0.06), as well as significant differences in PPA (t_(14)_ = 2.56, p = 0.01), MPA (t_(14)_ = 2.42, = 0.01), and EVC (t_(15)_ = 1.76, p = 0.049) (Supplemental Figure 3). However, repeated measures ANOVAs with factors for region (OPA, PPA, and MPA; or OPA and EVC) and condition (within, between) failed to reveal significant region x condition interactions (both F’s < 0.22, both p’s > 0.70).

**Supplemental Figure 3.**
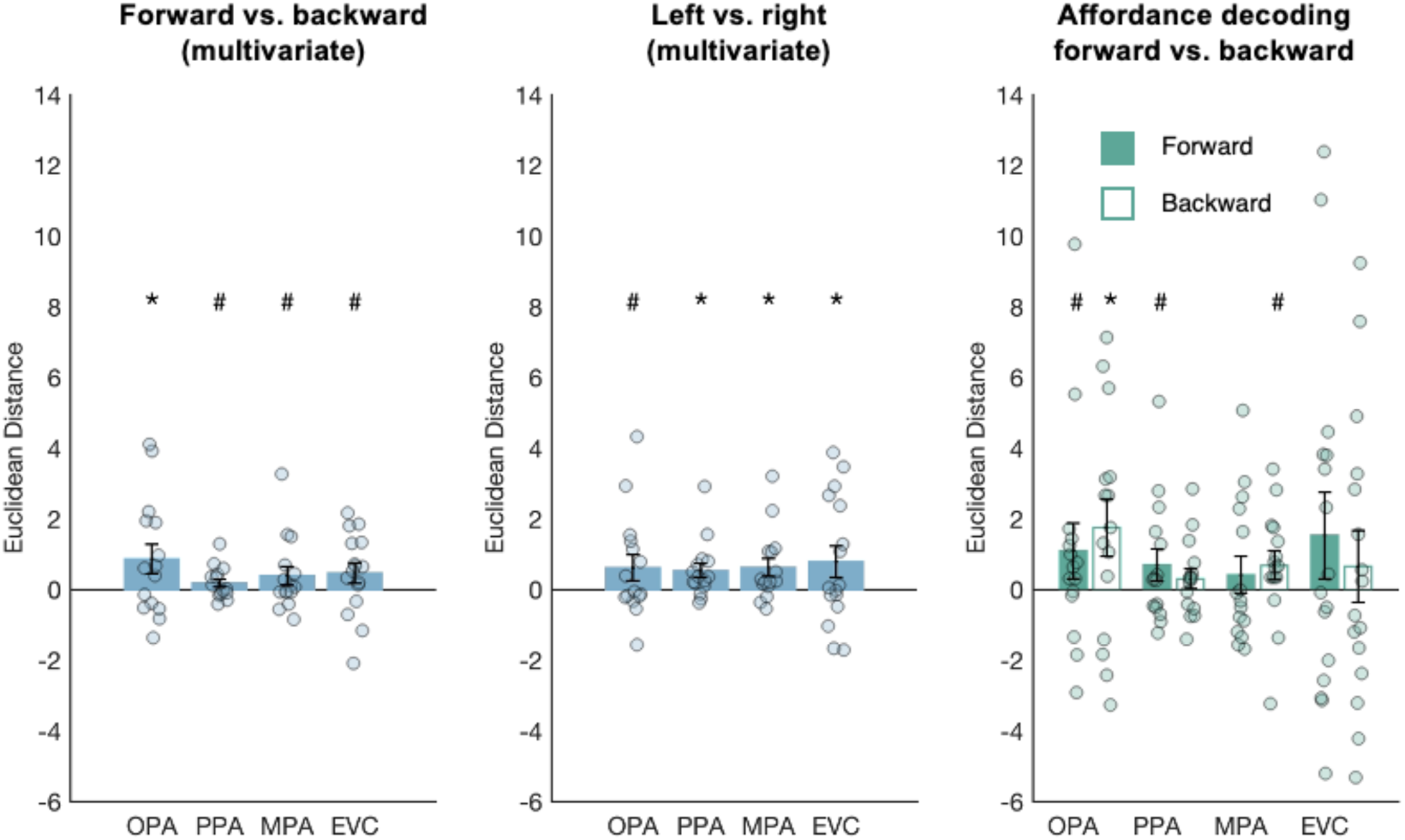
Additional tests of ego-motion representation. Bar plots indicate the difference score for the “between” conditions minus the “within” conditions; responses greater than zero indicate more similar responses (i.e., smaller Euclidean distance) for the “within” than “between” condition pairs, and thus significant decoding. Error bars represent the standard error of the mean. Markers indicate data from individual participants.

Finally, we asked: how are processing of navigational dynamics and structure related? Navigational dynamics might influence processing of navigational structure, since the structure of navigable space is more relevant in the direction of the current heading (e.g., ahead of you while walking forward) than in another direction (e.g., behind you while walking forward). To test this possibility, we analyzed conditions 1-6, which depicted forward or backward ego-motion through scenes with a door on either the left, right, or both walls. If navigational dynamics influence representation of navigational structure, then we will find stronger decoding of affordance direction (i.e., calculated as within-minus between-correlations for the left, right, and both conditions) for the forward conditions than the backward conditions, tested with a paired samples t-test (one tailed). We failed to find evidence of this effect in all four regions (all t’s < 1.31, all p’s > 0.10) (Supplemental Figure 3).

### Additional preregistered analyses of the integration of navigational affordances and ego-motion

If OPA integrates navigational affordances and ego-motion then OPA responses will differ depending on whether the depicted ego-motion is consistent with the navigational affordance of the space (e.g., a left turn toward a door on the left). To test this prediction, we analyzed conditions 7, 8, 9, and 10, which depicted left or right turns in the direction of either an open doorway that affords further navigation (i.e., a “consistent” turn) or away from a doorway toward a wall containing a painting, which does not afford further navigation (i.e., an “inconsistent” turn). Key analyses testing this prediction are already present in the primary results. For an additional univariate analyses, we predicted an overall difference in response to the consistent and inconsistent conditions, tested using a two-tailed paired samples *t*-test. We found a marginally greater response to expected than unexpected turns in OPA (t_(14)_ = 2.09, p = 0.06) and MPA (t_(14)_ = 1.91, p = 0.08), as well as a significantly greater response to expected than unexpected turns in PPA (t_(14)_ = 4.24, p < 0.001), and a significantly weaker response to expected than unexpected turns in EVC (t_(15)_ = −2.28, p = 0.04). Directly comparing between the three scene regions, a 3 (region: OPA, PPA, MPA) x 2 (condition: expected, unexpected) repeated measures ANOVA failed to reveal a significant region x condition interaction (F_(1.25, 16.25)_ = 1.46, p = 0.25). However, comparing OPA and EVC, a 2 (region: OPA, EVC) x 2 (condition: expected, unexpected) repeated measures ANOVA did reveal a significant region x condition interaction (F_(1, 14)_ = 16.57, p = 0.001).

